# Aging impacts basic auditory and timing processes

**DOI:** 10.1101/2024.03.24.586049

**Authors:** Antonio Criscuolo, Michael Schwartze, Leonardo Bonetti, Sonja A. Kotz

**Affiliations:** Department of Neuropsychology & Psychopharmacology, Faculty of Psychology and Neuroscience, Maastricht University, 6200 MD, Maastricht, the Netherlands; Center for Music in the Brain, Department of Clinical Medicine, Aarhus University & The Royal Academy of Music, Aarhus, Aalborg, Denmark; Centre for Eudaimonia and Human Flourishing, Linacre College, University of Oxford, Oxford, United Kingdom; Department of Psychiatry, University of Oxford, Oxford, United Kingdom; Department of Neuropsychology, Max Planck Institute for Human Cognitive and Brain Sciences, 04103, Leipzig, Germany

**Keywords:** aging, timing, audition, oscillations, EEG

## Abstract

Deterioration in the peripheral and central auditory systems is common in older adults and often leads to hearing and speech comprehension difficulties. Even when hearing remains intact, electrophysiological data of older adults frequently exhibit altered neural responses along the auditory pathway, reflected in variability in phase alignment of neural activity to speech sound onsets. However, it remains unclear whether these challenges in speech processing in aging stem from more fundamental deficits in auditory and timing processes. Here, we investigated *if* and *how* aging individuals encoded temporal regularities in isochronous auditory sequences presented at 1.5Hz, and *if* they employed adaptive mechanisms of neural phase alignment in anticipation of next sound onsets. We recorded EEG in older and young individuals listening to simple isochronous tone sequences. We show that aging individuals displayed increased amplitudes and variability in time-locked responses to sounds, an increased 1/F slope, but reduced phase-coherence in the delta and theta frequency-bands. These observations suggest a lack of repetition-suppression and inhibition when processing repeated and predictable sounds in a sequence and altered mechanisms of continuous phase-alignment to expected sound onsets in aging. Given that deteriorations in these basic timing capacities may affect other higher-order cognitive processes (e.g., attention, perception, and action), these results underscore the need for future research examining the link between basic timing abilities and general cognition across the lifespan.

**Highligths:**

- Aging individuals (HO) show increased cortical excitability as compared to younger adults (HY);
- HO’s neural responses to fully predictable isochronous tones were larger and more variable than HY;
- HO showed reduced phase coherence in delta- and theta-band oscillations during listening to auditory sequence;
- Altogether, these results show altered sensory and timing processes in aging.

## 1. Introduction

### ‘Wait, what? Can you repeat it?’

A cascade of biochemical, neuro-functional and -anatomical changes takes place in aging. Deteriorations in the peripheral (e.g., loss of hair, ganglion and/or striatal cells) and central auditory systems [1,2] are particularly common and typically lead to a decline in auditory processing capacity [3–6]. However, structural brain changes often extend more broadly, and include widespread reductions in grey and white matter volume across the brain [7], as well as in cortico-subcortical connectivity [8]. Moreover, modifications within striatal-frontal networks [9], under-recruitment of the cerebellum during challenging cognitive tasks [10], and alterations in cerebellum-basal ganglia connectivity [11] have been linked to diminished cognitive control [12] and a variety of motor and cognitive deficits [11]. However, there is significant heterogeneity in the trajectories of neurocognitive and structural decline, stemming from substantial inter-individual variability in risk and modulating factors [13]. Furthermore, there exists variability in the capacity to compensate for cognitive decline by recruiting additional neural resources and/or adopting compensatory cognitive strategies [13]. For example, despite inevitable hearing loss [1,2], speech comprehension is largely preserved in older adults [14,15]. Performance, however, declines rapidly in challenging listening conditions, and is accompanied by decreased activation of the auditory cortex [15], inferior frontal regions, and reduced connectivity within the speech network [14]. Aging individuals tend to engage more working memory and attentional networks (e.g., frontal and prefrontal regions) in a compensatory manner [15]. Even in the absence of hearing loss, evidence confirms general difficulties in encoding simple and complex sounds, beginning in the brainstem [16] and in the inferior colliculus [17]. Auditory nerve modeling has demonstrated that the deterioration of auditory nerve fibers and loss of inner hair cells impact the brain’s capacity to precisely phase-lock (i.e., align) neural responses to sound onsets [18], resulting in reduced amplitude and phase-coherence of brainstem responses to simple and complex sounds [16]. This, in turn, can affect speech processing, as indicated by variable brainstem responses, decreased phase-locking to speech sounds [6,19], and a reduced connectivity between the brainstem and auditory cortex [4]. The weakened sensitivity to auditory input via the brainstem is typically compensated by increased excitability of the auditory cortex [20] and altered responses to sounds [3,6,21–24]. Consequently, event-related potentials (ERPs) recorded by electroencephalography (EEG) exhibit enhanced amplitude responses in aging individuals, particularly in the N100 component [3,20,21,23,25–27]. There is consensus in associating these larger ERP responses with the reduced ability to employ ‘sensory gating’ [26], an adaptive mechanism to suppress cortical responses to repetitions of predictable stimuli [21,27]. Furthermore, variability in the latency of event-related responses to sounds [25,28] and the reduction in steady-state responses to auditory metronomes [29,30] suggest deteriorations in the encoding of the precise timing of sensory events, and in internalizing temporal regularity in auditory sequences. At the same time, there is complementary evidence showing greater neural synchronization with amplitude and frequency modulations of sounds in aging individuals and increased sensitivity to temporal regularities, as revealed by metrics of phase concentration [17,22,24,31]. Larger and less variable event-related responses [23] were, however, typically accompanied by reduced sustained neural activity to sound modulations in continuous listening scenario [22,24,32]. These partially contradicting observations leave open the question of whether aging impacts the *basic* capacities to *detect* temporal regularities in the sensory environment, generate predictions about the *timing* of future events, and employ these predictions to optimize sensory processing and perception. In turn, this perspective prompts the question: are difficulties in speech comprehension observed in older adults linked to speech-specific processing difficulties or to more fundamental temporal processing deficits?

We aimed to investigate whether older adults detect, encode, and employ temporal regularities in the sensory environment to generate predictions and optimize sensory processing similarly to younger adults. We addressed this question by recording EEG in older and younger adults while they listened to isochronous tone sequences and performed a deviant counting task. The task served the scope to focus their attention on the formal properties of the auditory sequences, while diverting their attention from temporal regularity. As such, we did not directly instruct participants to process the timing of sound onsets. We hypothesized that aging would be associated with increased variability in event-related responses to tone onsets, as indexed by metrics of N100 variability, and reduced Inter-Trial Phase Coherence (ITPC) in delta-band oscillatory activity. Furthermore, we hypothesized that older individuals would fail to show a continuous phase-alignment while listening to isochronous sequences. Finally, we expected a steeper 1/F of the Fourier spectrum in the aging group, indicative of increased cortical excitability. Combined results from event-related, spectral parametrization, and ITPC analyses confirmed increased cortical excitability and hypersensitivity to sound onsets, and reduced phase coherence in delta- and theta-frequency bands in the aging group. Altogether, these observations suggest that aging alters basic sensory and temporal processing.

These limitations in fundamental timing capacities in older adults might critically affect not just basic auditory processing but also higher-order cognitive functions such as speech processing. Consequently, the current results motivate future research on the impact of altered timing capacities on cognition in aging and across the lifespan.

## 2. Materials & Methods

### 2.1. Participants

Forty-three native German speakers participated in this study and signed written informed consent in accordance with the guidelines of the ethics committee of the University of Leipzig and the declaration of Helsinki. Participants were grouped into 18 younger (HY; 9 females; 21–29 years of age, mean 26.2 years) and 18 older (HO; 9 females; 50-78 years of age, mean 60 years) adults. All participants were right-handed, had normal or corrected-to-normal vision, and no hearing deficits. Participants received 8€/h for taking part in the study. Participants were not asked to indicate musical expertise and/or daily music listening choices.

### 2.2. Experimental design and procedure

Participants listened to 96 sequences comprising 13-to-16 tones (F0 = 400Hz, duration = 50ms, amplitude = 70dB SPL; standard STD), presented in one recording session of approximately 25min. Each tone sequence included one or two deviant tones (DEV), attenuated by 4dB relative to the STD tones. The inter-onset-interval between successive tones was 650ms, resulting in a stimulation frequency (*Sf*) of 1.54Hz, and a total sequence duration of 8.45-10.4s (13 to 16 tones * 650ms; Fig. 1A). Participants were seated in a dimly lit soundproof chamber facing a computer screen. Every trial started with a fixation cross (500ms), followed by an auditory sequence. The cross was continuously displayed on the screen to prevent excessive eye movements while listening to the auditory sequences. At the end of each sequence, a response screen appeared and prompted participants to immediately press a response button to indicate whether they had heard one or two softer tones. After the response, there was an inter-trial interval of 2000ms. A session was divided into two blocks of approximately 10 minutes each, with a short pause in between (about 25min total duration). Stimulation materials and experimental setups thus mirror those adopted and previously described [33–35].

**Figure 1.**
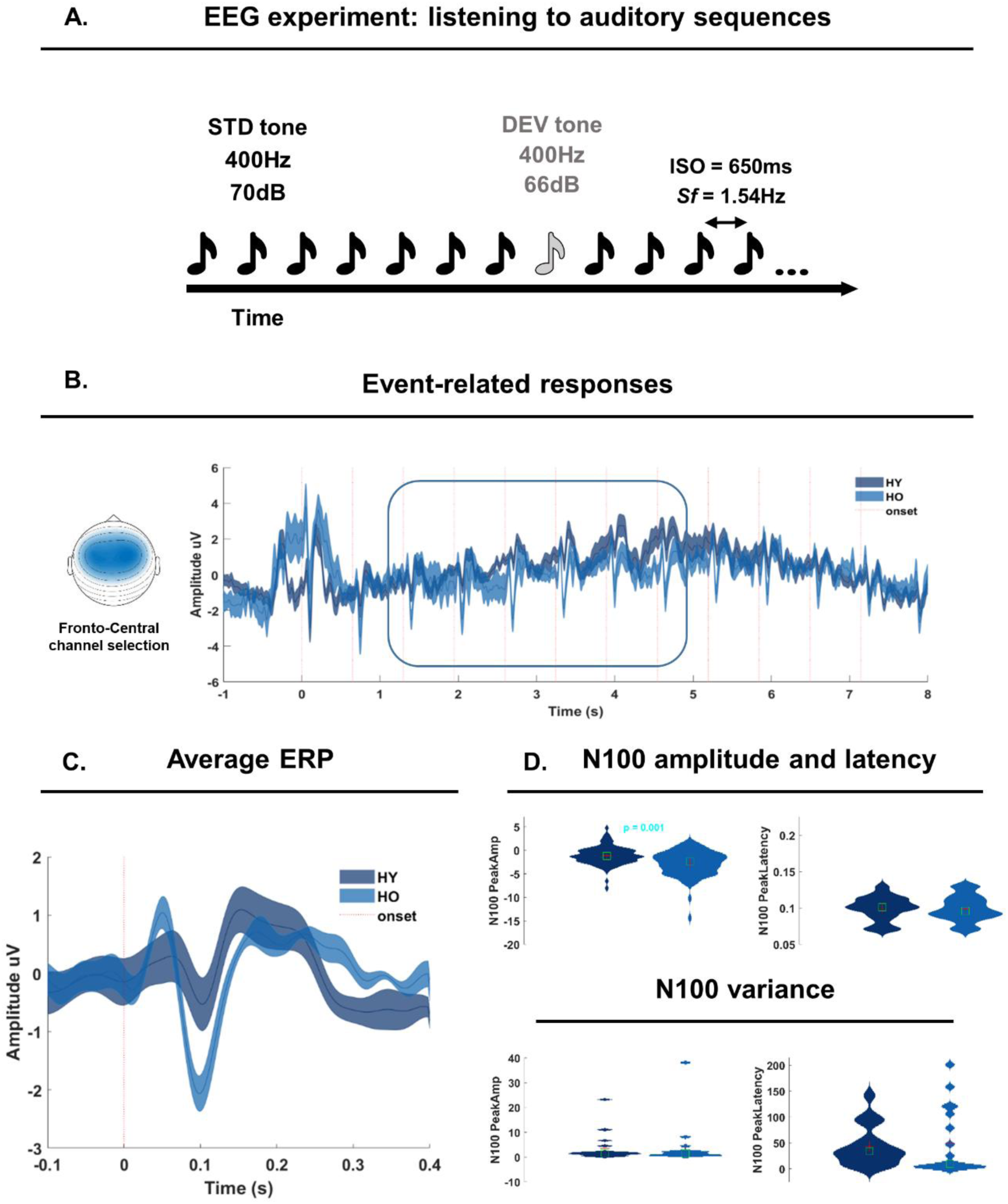
- Experimental setup and Event-related analyses. A. Participants listened to 96 isochronous tone sequences, containing 13-to-16 tones including standard (STD) tones (F0 = 400 Hz, duration = 50 ms, amplitude = 70dB SPL) and either one or two deviants (DEV; attenuated by 4dB relative to the STD tones). The first DEV could either fall on positions 8,9,10, or 11, while the second DEV always fell on position 12. The inter-onset-interval between successive tones was 650ms, resulting in a stimulation frequency (Sf) of 1.54Hz. B. Event-related analyses focused on a fronto-central (FC) channel cluster (as provided on the left side) and on 5 tones from the 3rd to the 7th position along the auditory sequence (square on the time-series). Dark blue lines report the time-series for younger (HY) participants, while the lighter blue lines report the time-series for older adults (HO) participants. C. Average ERPs over 5 tone positions as highlighted in B, and in the FC channel cluster. Color coding as in B. D. N100 peak amplitude and latency (top panel) were statistically compared across the two groups by means of permutation testing. At the bottom, the variance in the N100 peak amplitude and latency was statistically compared across the two groups by means of permutation testing.

### 2.3. EEG recording

The EEG was recorded from 59 Ag/AgCl scalp electrodes (Electrocap International), amplified using a PORTI-32/MREFA amplifier (DC set to 135Hz), and digitized at 500Hz. Electrode impedances were kept below 5kΩ. The left mastoid served as an online reference. Additional vertical and horizontal electro-oculograms (EOGs) were recorded.

### 2.4. Data Analysis

#### 2.4.1. EEG Preprocessing

The preprocessing pipeline and the analysis approach adopted here mirror and expand those described previously [33–35]. EEG data were analyzed in MATLAB with a combination of custom scripts and functions and the FieldTrip toolbox [36]. Data were first re-referenced to the average of the two mastoid electrodes and band-pass filtered with a 4th order Butterworth filter in the frequency range of 0.1-50 Hz (*ft_preprocessing*). Eye-blinks and other artifacts were identified using independent component analysis. This semi-automated routine combined two steps: in the first iteration, we employed ‘*fastICA’* (as implemented in FieldTrip) to decompose the original EEG signal into independent components (N= number of EEG channels −1), then automatically identified components with a strong correlation (>.4; labeled as ‘bad’ components) with the EOG time-courses, removed them with ‘*ft_rejectcomponent*’, and then reconstructed the EEG time-course. In a second step, we again used ‘*fastICA’* but now with a dimensionality reduction to 20 components. We visually inspected these components via ‘ft_*rejectvisual’,* and selected ‘outliers’ (e.g., based on max values and z-scores). The 20 components were visually inspected after plotting their topographies and time-series, and a new selection of ‘outliers’ was defined. Lastly, we visually inspected the two lists of outliers and decided which components had to be removed. On average, we removed 2 components via ‘*ft_rejectcomponent*’. Then EEG time-series were reconstructed. In the next preprocessing step, we performed artifact subspace reconstruction as implemented in the ‘*pop_clean_rawdata*’ function in EEGlab, and with the ‘BurstCriterion’ parameter set to 20 (all other parameters were set to ‘off’). We then employed an automatic channel rejection procedure to remove noisy channels. In this routine, we calculated the median variance across channels (and excluding EOG channels), and ‘outliers’ were then defined as exceeding 2.5*median variance. Next, we implemented an artifact suppression procedure [33–35], a cleaning routine that interpolates noisy (>absolute mean+4*SD) time-windows on a channel-by-channel basis. Lastly, data were low-pass filtered at 40Hz via ‘*ft_preprocessing*’, segmented to each auditory sequence (starting 4s before the first tone onset and ending 4s after the last tone onset), and downsampled to 250Hz.

#### 2.4.2. Event-related analyses

We assessed the amplitude, latency, and variability of neural responses to tone onsets along the auditory sequences by adopting an event-related potential (ERP) approach. Thus, sequence-level data as obtained from preprocessing were further segmented into time-windows ranging from −1 to 8s relative to the first tone onset in each auditory sequence and later underwent a low-pass filter with a 20Hz frequency cutoff (‘ft_preprocessing’). Next, we centered the data by mean correcting each trial by a global average (calculated from −1 to 8s and across trials) and performed ‘peak analyses’. Thus, we calculated the participant-, trial-, and channel-level peak amplitude, latency and variability of the N100 component of the ERP [3,20,21,23,25–27]. For doing so, we defined a 60ms-long time-windows centered at 100ms. Within this time-window we obtained the amplitude peak and its latency (the max value and its time point). Next, we calculated the intra-individual variability (peak amplitude and latency) across trials and within a fronto-central channel (FC) cluster of interest. The FC cluster encompassed the sensor-level correspondents of prefrontal, pre-, para-, and post-central regions highlighted in previous studies [37] and further highlighted in similar EEG work on rhythm processing [33,34,38]. The cluster included 16 channels: ‘AFz’, ‘AF3’, ‘AF4’, ‘F3’, ‘F4’, ‘F5’, ‘F6’, ‘FCz’, ‘FC3’, ‘FC4’, ‘FC5’, ‘FC6’, ‘C1’, ‘C2’, ‘C3’, ‘C4’. As the first tones within an auditory sequence are known to elicit much stronger neural responses compared to later tones, we focused subsequent analyses on tones from the 3^rd^ to the 7^th^ position (STD before the onset of a DEV tone).

##### Statistical analyses

Statistical analyses assessed group differences in the N100 peak amplitude over tone repetitions along the auditory sequence by means of a repeated-measure ANOVA. Thus, individual N100 peak amplitudes over 5 tonal positions (3^rd^ to 7^th^) were modelled by the ‘fitrm’ algorithm by specifying ‘Group’ and ‘Time’ as factors and allowing for an interaction term. Next, the model entered a repeated measures analysis of variance via the ‘ranova’ function. In the absence of a Group x Time interaction, we proceeded by testing for the main effect of Group.

##### Mixed Effect Models on ERP data

We assessed group differences in N100 Peak amplitude and latency via Mixed effect models. The model included ‘Group’ as a fixed factor and a random intercept per participant. Model information and results are reported in Tab 1 and Suppl. Tab. 1, respectively.

##### Statistical comparisons on the variance

We assessed group differences in the variability of N100 Peak amplitude and latency by means of permutation testing. This iterative procedure performs 1000 permutations of data points belonging to one or the other group and ultimately assess the p value from the original groups against the p obtained from permutations. A *p-*value lower than .05 was considered statistically significant. Results are provided in Fig. 1D.

#### 2.4.3. Spectral parametrization

To investigate how participants (i) encoded temporal regularities in auditory sequences, (ii-iii), and whether there were group differences in the excitation/inhibition balance [39,40], we performed spectral parametrization analyses. Differently from typical Fourier (FFT) analyses, the spectral parametrization allows disentangling oscillatory from non-oscillatory components (i.e., the 1/F typically observed in the Fourier spectra) [41]. Thus, a series of tone-locked neural responses should lead to a clear amplitude peak in the frequency spectrum at the *Sf.* To test this hypothesis, we first shortened trials into segments of 8s (from the first tone onset (0s) to the 12^th^ tone offset), and then employed the automated spectral parameterization algorithm described in [41] and implemented in FieldTrip in a two-step approach. Thus, ‘ft_freqanalysis’ was first used in combination with the multi-taper method for FFT (‘mtmfft’) and power as output (‘pow’), and secondly by specific ‘fooof_aperiodic’ as output. The output frequency resolution was set at 0.2Hz. Next, the aperiodic (fractal) spectrum was removed from the FFT spectrum via calling the ‘ft_math’ function, finally isolating the so-called ‘oscillatory’ (Fig. 2A, left) from a non-oscillatory (‘fractal’) component (Fig. 2A, right).

**Figure 2.**
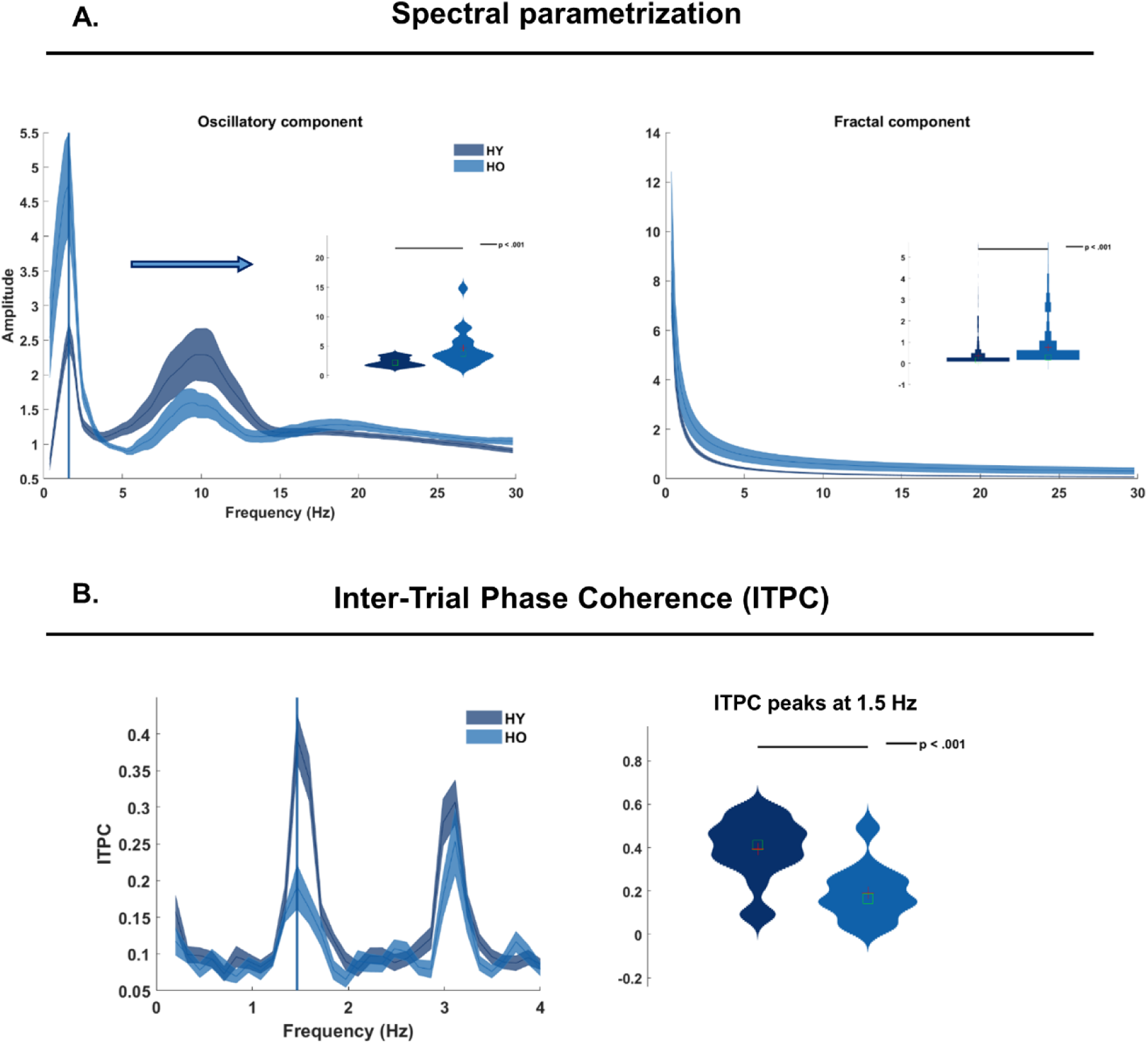
– Spectral parametrization and Inter-trial phase coherence analyses. *A.* Spectral parametrization analyses allowed to decompose the Fourier spectrum into an oscillatory component (left) and into a non-oscillatory (fractal) component (right). Both spectra provide frequencies on the x-axis and amplitude values on the y-axis. Dark blue lines report the data for younger (HY) participants, while the lighter blue lines report the data for older (HO) participants. The inserts on top of the main panel provide the statistical comparisons on individual data as assessed by permutation testing. On the left, the group comparison assessed the amplitude peak at the stimulation frequency (1.5Hz). On the right, the group comparison assessed for amplitude differences across the entire spectrum. *B.* Inter-trial phase coherence (ITPC) analyses assessed group differences in the ITPC at the stimulation frequency by means of permutation testing (right). The ITPC spectrum (left) provides frequencies on the x-axis and coherence values on the y-axis. The two vertical lines on the spectrum report the group coherence peak. *C.* Exploratory analyses assessed the link between peak amplitudes in the oscillatory component (on the left, FooFosc) and in the ITPC (right) with ERP metrics of peak amplitude (Amp), latency (Lat) and variance (Var) across three ERP components (P50, N100, P200). Dark blue lines report positive relations, while lighter blue lines report negative relations.

##### Statistical analyses

Subsequent statistical analyses were performed on the same FC cluster as described above. Group differences were statistically assessed by permutation testing (1000 permutations) of the extracted peak amplitude values at the *Sf* and the amplitude of the fractal component across the frequency spectrum. A *p-*value below .05 was considered statistically significant.

#### 2.4.4. Inter-Trial Phase Coherence

When neural activity precisely encodes the temporal regularities in auditory sequences, it should not only show a clear amplitude peak in the FFT spectrum but also display phase coherence. The inter-trial phase coherence (ITPC) metric is inversely proportional to the variability in the imaginary part of the complex FFT spectrum. Thus, when oscillations are precisely aligned over trials (they have the same phase), the ITPC is high; when, instead, there is variability in the phase of the oscillations over trials, the ITPC is lower.

The complex FFT spectrum was obtained by performing FFT decomposition at the single-participant, -channel and -trial level on 8s-long segments as above. Next, the ITPC spectrum was calculated by dividing the Fourier coefficients by their absolute values (thus, normalizing the values to be on the unit circle), calculating the mean of these values, and finally taking the absolute value of the complex mean. Further documentation can be found on the FieldTrip website (https://www.fieldtriptoolbox.org/faq/itc/). For illustration purposes, the ITPC spectrum was restricted to 1-4Hz (Fig. 2B).

##### Statistical analyses

Subsequent statistical analyses were performed on the same FC cluster as for the event-related analyses. Group differences were statistically assessed by permutation testing of the extracted ITPC values at the *Sf* and with 1000 permutations. A *p-*value below .05 was considered statistically significant.

##### Time-resolved ITPC

Although ITPC and similar phase concentration measures are typically interpreted as a proxy of *entrainment*, they mostly reflect a sequela of time-locked evoked responses [42–45]. In turn, the variability in the latency of event-related responses is inversely proportional to the estimated ITPC. As ITPC mostly depends on the (initial) phase estimates from fast-Fourier transformations, it typically does not allow assessing the dynamics of phase alignment in a continuous manner. In other words, we cannot test whether phase coherence changes during the listening period.

Here, we estimated a time-resolved metric of ITPC (t-ITPC) to quantify the build-up of phase-coherence over the course of an auditory sequence. For doing so, we first employed time-frequency transform (TF data), then calculated the t-ITPC using the complex spectra of TF data (same procedure as in the ITPC above) and lastly calculated the slope of the t-ITPC at the single-participant level and across frequency-bands. More details are provided in the respective paragraphs below.

#### Time-frequency transform

After preprocessing, single-trial EEG data underwent time-frequency transformation (‘*ft_freqanalysis’*) by means of a wavelet-transform [46]. The bandwidth of interest was centered around the stimulation frequency (+/- 1Hz, i.e., .54 - 2.54Hz, thus obtaining a 1.54Hz center frequency), using a frequency resolution of .2Hz. The number of fitted cycles was set to 3. The single-trial approach resulted in ‘induced’ (as compared to ‘evoked’) responses. The output was a complex spectrum; no averaging over channels, trials, or participants was performed at this stage.

#### Slope of t-ITPC

As for the ITPC above, t-ITPC was obtained by dividing the complex coefficients of TF-data by their absolute values (thus, normalizing the values to be on the unit circle), averaging, and finally taking the absolute value of the complex mean (average over participants in Fig. 3A). Next, we calculated the slope of each t-ITPC time-series (e.g., Fig. 3B) by fitting a first-order polynomial (‘*polyfit*’ function in MATLAB; *p*) and then deriving a first-order approximation (*p(1)*Time+p(2)*). The calculation of the slope was performed by starting from the 3^rd^ tone onset and up to the 8^th^ tone (where the DEV tone was likely to occur).

**Figure 3.**
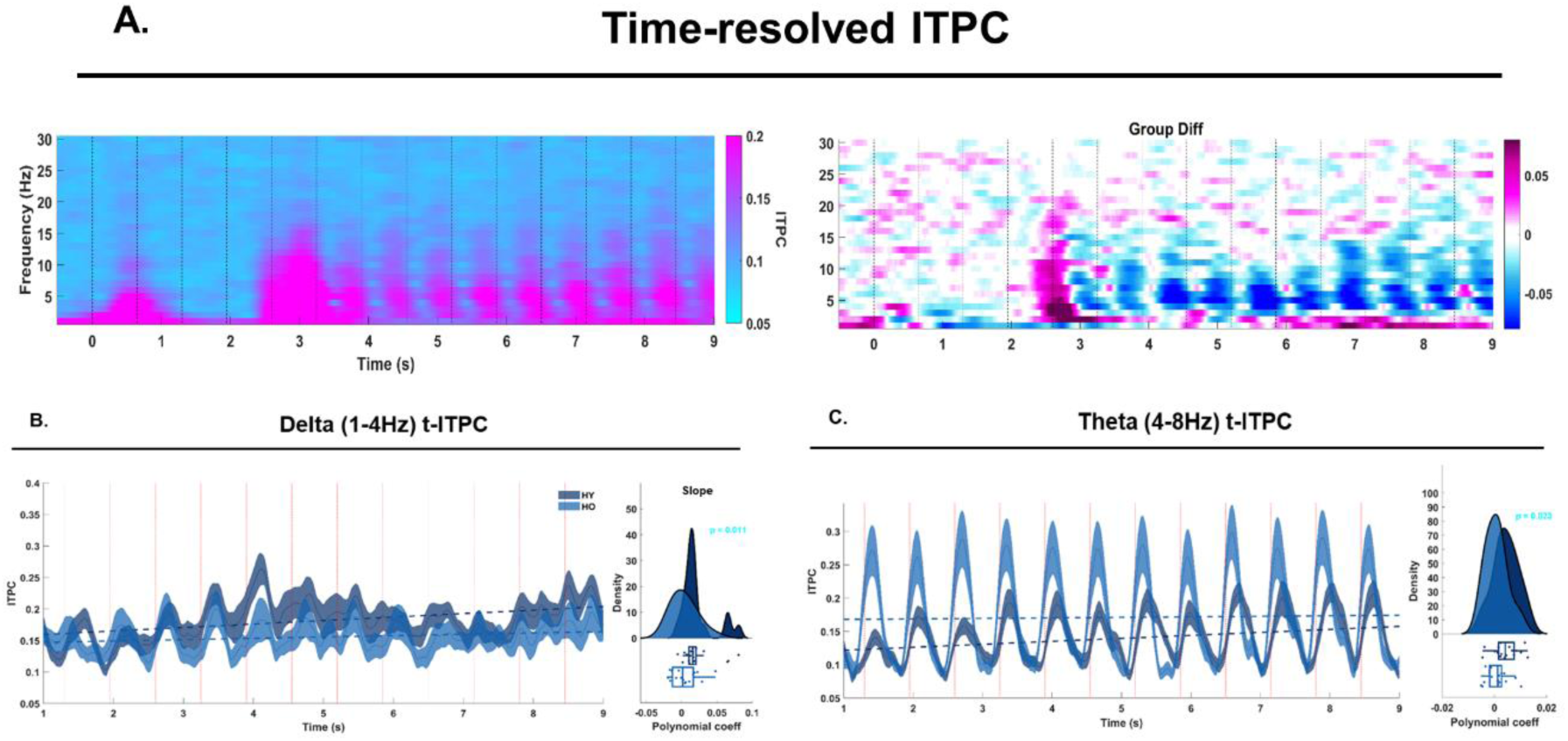
– Time-resolved ITPC. A, left: the time-course of time-resolved inter-trial phase coherence (t-ITPC) time-locked to the auditory sequence and averaged across all participants. The time interval shown in the x-axis ranges from −1 to 9s. Vertical dotted lines indicate tone onsets. On the y-axis frequency in Hz, ranging from 1 to 30Hz. The color bar reports low ITPC values in light blue and higher ITPC values in pink. A, right: the time-course of t-ITPC differences comparing HY and HO (HY minus HO). The x- and y-axis are the same as in the left panel. The color bar reports greater ITPC values in the HO (>HY) in blue, and greater ITPC values in the HY (>HO) in pink. B: the time-course of the delta-band t-ITPC for HY (darker blue) and HO (lighter blue). Shaded contours report the standard error calculated per group across participants. The vertical dotted lines represent tone onsets. The horizontal dotted lines report the slope of the t-ITPC per group. On the right, the distribution of the slope coefficient across participants, per group. C: the time-course of theta-band t-ITPC for HY and HO as in panel C.

##### Statistical comparisons

Group differences in the slope of the t-ITPC at each frequency-band were assessed via permutation testing and with a total of 1000 permutations.

### Data and code Availability

The analysis code and data in use here can be provided upon reasonable request by the corresponding author.

## 3. Results

### 3.1. Event-Related Analyses

Event-related analyses tested for group differences in the N100 component of the event-related potential (ERP; Fig. 2B-D). A repeated-measures ANOVA tested for group differences in the N100 peak amplitude over 5 tone positions along the auditory sequences (3^rd^ to 7^th^ position). This analysis specifically assessed a repetition-suppression effect. The Group * Time interaction term of the model was not significant (*F* = .63, *p* = .49). We then tested the main effect of Group across the sequence, by pooling the N100 peak amplitudes across the 5 tone positions. The group effect was statistically assessed by mixed effect models and including a fixed effect of group and a random intercept per participant. The model reported a significant group difference in the N100 peak amplitude (t (2,180) = −3.24, p = .001; Tab 1). The mixed effect model testing group differences in the N100 peak amplitude latency was not significant (Suppl. Tab. 1).

Next, we assessed group differences in the variability of the N100 peak amplitude and latency. Neither of the two comparisons reported significant group differences.

**Table 1.**
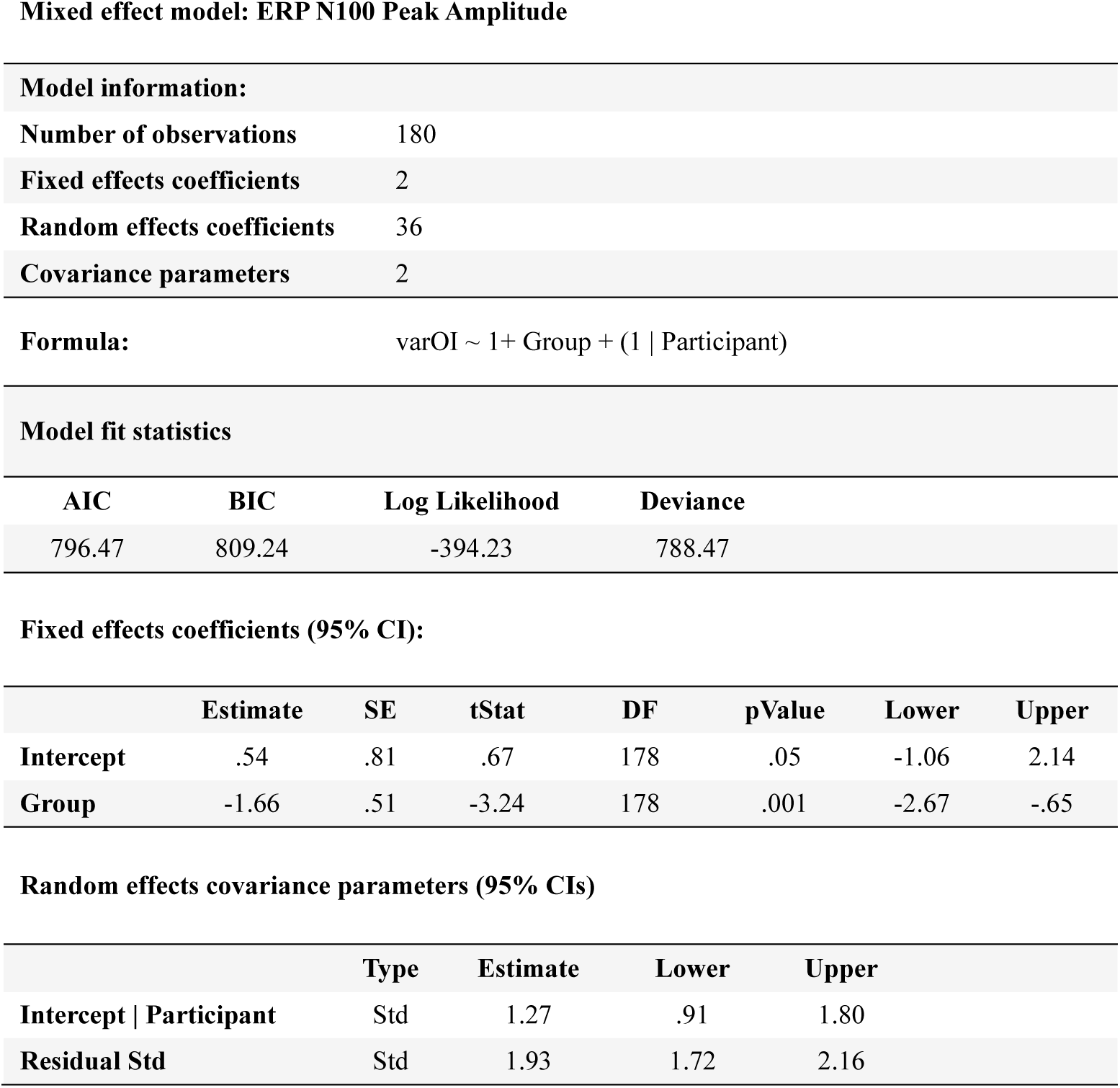
- Mixed effect model on ERP N100 Peak Amplitude.

The table reports model information: number of observations, fixed effect coefficients, random effect coefficients, covariance parameters. Then, the formula used to fit the model and model fit statistics: AIC, BIC values, Log Likelihood and Deviance. Further below, the fixed effect coefficients in a 95% confidence interval (CI): estimate, standard error, t-stat, degrees of freedom (DF), p value, lower and upper bound. Right below, random effects covariance parameters: estimate, lower and upper bound.

### 3.2. Spectral parametrization

After decomposing the Fourier spectrum into a so-called ‘oscillatory’ component (OSc) and a fractal component (FRc), we statistically assessed group differences in the amplitude of the OSc at the stimulation frequency (1.5Hz; *Sf*) and the amplitude of the FRc across frequencies by means of permutation testing, and with 1000 permutations. The group effect for the OSc at the *Sf* was statistically significant and showed larger amplitude responses in the HO than the HY (*p* < .001; Fig. 2A, left). HO also showed a significantly stronger FRc across the spectrum (*p* < .001; Fig. 2A, right).

### 3.3. Inter-Trial Phase Coherence

The imaginary part of the complex Fourier spectrum was used to calculate the Inter-trial phase coherence (ITPC). Statistical analyses assessed group differences in ITPC at the *Sf*. Permutation testing revealed a significant group effect, with HY showing larger ITPC at the *Sf* as compared to HO (*p* = .001; Fig. 3B).

### 3.4. Time-resolved ITPC

T-ITPC was obtained by quantifying phase coherence from the complex spectra of continuous wavelet transformed data in the delta-, theta-, alpha- and beta-frequency bands. The group average t-ITPC time course is displayed in Fig. 3A, left. The group difference across frequency-bands is provided in Fig. 3A, right. For each participant and frequency band, we calculated the slope of t-ITPC and later performed group comparisons via permutation testing. HO showed a reduced t-ITPC slope in the delta (Fig. 3B) and theta (Fig. 3C) frequency-bands, but not in the alpha and beta-bands.

## 4. Discussion

Throughout the lifespan, neuroanatomical brain changes typically follow an inverted U-shape trajectory [47]. Grey and white matter volume increase from childhood to adulthood and then decline with aging. Concurrently, primary sensory systems often undergo gradual deterioration, potentially leading to decline in auditory processing [3,5,20]. Changes within both peripheral and central auditory systems, such as the deterioration of auditory nerve fibers and loss of inner hair cells, affect the brain’s ability to accurately encode sensory events in the auditory environment [18], consequently impacting speech comprehension, social interactions, and cognition more broadly [48]. Importantly, even in the absence of hearing loss, older individuals experience difficulties in processing both simple and complex auditory sequences in noisy environments [16,25,26,28].

In this study, we posited that the challenges observed in higher cognitive processes such as speech processing during aging might stem form an underlying decline in the ability to detect, encode, and employ temporal regularities in the sensory environment to optimize sensory processing. To investigate this hypothesis, we recruited younger and older individuals and recorded their neural activity using EEG while they listened to simple isochronous equitone sequences presented at a stimulation frequency (*Sf*) of 1.5Hz. Analyses of event-related potentials (ERP) and spectral data (spectral parametrization analyses) revealed greater evoked responses in older adults, consistent with previous findings documenting hypersensitivity to sensory input in aging [3,20,32,21–27,31]. These results support the notion that aging affects the ability to engage in ‘sensory gating’ [26], thus failing to adaptively inhibit or suppress cortical responses to repetitive and predictable stimuli [21,24,27,49].

Spectral parametrization analyses further showed that older adults exhibited an increase in the amplitude of the (1/F) fractal component across frequencies. This observation is consistent with the enhanced event-related responses described earlier and supports the notion of heightened excitability (or reduced inhibition) of the auditory cortex in aging [20]. While prior evidence showed diminished encoding of temporal regularity in metronome-like auditory sequences (obtained through typical Fourier analyses) [29,30], the removal of the fractal component from the frequency spectrum allowed revealing the reversed pattern: enhanced evoked responses to sounds in older individuals. This result seems to confirm prior evidence showing greater neural synchronization with sounds modulations in aging individuals, and an increased sensitivity to temporal regularities [17,22,24,31]. However, the inter-trial phase coherence (ITPC) analyses showed that older adults exhibited lower ITPC at the stimulation frequency (*Sf*; 1.5Hz), ultimately indicating increased variability in the neural encoding of sound onsets. Thus, we showed that aging is associated with stronger evoked responses, but reduced phase-alignment. These results support previous observations of reduced coherence and phase alignment of neural activity to sounds in simple and more complex auditory sequences [6,16,18,19].

The hypersensitivity observed in aging individuals is typically further accompanied by reduced sustained neural activity in continuous listening scenario [22,24,32], potentially affecting speech tracking and comprehension, especially in noisy environments [31,32]. In order words, while older individuals display larger cortical responses to sound onsets, they seem to have difficulties in tracking the temporal fluctuations in the speech envelope [31].

We here asked whether aging individuals would show similar difficulties in tracking and anticipating simple sound onsets in isochronous contexts. To assess the continuous neural dynamics of onset tracking, we employed a time-resolved ITPC coherence metric and calculated the slope of phase coherence during continuous listening. We showed a flatter slope of t-ITPC in the delta- and theta-band activity in aging individuals, but no differences in higher frequency bands (alpha and beta). This result confirms a reduction in sustained neural activity and tracking of sound onsets in the aging brain, and further confirms that alpha-band regulatory mechanisms are intact in older individuals [50].

Taken together, these observations suggest that aging impacts basic sensory and timing processes. These findings underscore the importance of future research investigating the relationship between basic timing capacities and higher-order cognitive processes (e.g., speech processing) across the lifespan.

This evidence, however, challenges the ‘exploration-exploitation shift’ hypothesis [12]. According to this perspective, most aging individuals attempt to counteract sensory and cognitive decline by adopting compensatory cognitive strategies and leveraging on previous knowledge predictively [51]. For example, they may utilize long-term knowledge, generalizations, and predictions to mitigate increased difficulty with learning and the decline of executive functions due to striatal cholinergic changes [52]. To operationalize and test this notion, Brown et al., [51] referred to the predictive coding framework: given the reduced certainty of sensory signals, aging individuals rely more on memory and consequently generate predictions about future events [53]. These predictions serve the purpose of adaptation aiming to optimize perception and cognition despite cognitive decline [54]. Contrary to these expectations, the current findings indicate that older individuals either did not form predictions, or, at the very least, did not utilize temporal predictions to inform sensory processing at the very fast, millisecond temporal scale. Indeed, consistent with prior electrophysiological evidence, neural responses were not attenuated by top-down modulatory suppression mechanisms [3,20,21,23,25–27].

The abilities to detect and encode temporal regularities, as well as to form temporal predictions, have been linked to widespread cortico-subcortical circuitries including the basal ganglia and the cerebellum [55]. Lesions in either of these circuitries have been shown to causally impact the ability to predictively align neural dynamics to sound onsets [33]. Conversely, aging is typically characterized by decreased fractional anisotropy and increased diffusivity, indicative of white matter deteriorations, along with bilateral grey [57] and white matter loss in the cerebellum (CE) and reduced connectivity within the dentato-thalamo-cortical network [10]. Additionally, functional connectivity patterns undergo alterations [58], such as reduced within-network connectivity and variegated patterns of increase and decrease in between-network connectivity [59]. Notably, deteriorations within the striatal-frontal networks [9], the under-recruitment of the cerebellum during challenging cognitive tasks [10], and changes within the cerebellum-basal ganglia circuitries [11] have been associated with reduced cognitive control [12] and numerous motor and cognitive deficits [11].

Aligned with the original hypotheses and bolstered by these novel findings, we propose that neuroanatomical and functional alterations in cortico-subcortical circuitries, including the basal ganglia (BG) and the cerebellum (CE) may impact fundamental timing and predictive abilities, which are integral to cognition. These changes could impact the documented declines in processing speed, working memory, inhibition, memory, and reasoning capacities [13,60,61]. However, establishing a causal link between timing, predictive functions, and general cognition presents a challenge due to the substantial heterogeneity in aging trajectories. Indeed, neuroanatomical and cognitive changes throughout the life are subject to modulation by a complex interplay of vascular, metabolic, and inflammatory risk factors [7], which, in turn, are influenced by the intricate interaction of environmental factors (e.g., socioeconomic status and education) and genetic predispositions [61]. Variability in any of these modulating variables inevitably results in significant diversity in cognitive capacities among older adults, impeding generalizations. Therefore, systematic, longitudinal, and comprehensive assessments of timing and cognitive functions across the lifespan are imperative.

## 5. Conclusions

Here, we examined the effects of aging on basic sensory and temporal processing. The integration of findings from three complementary analytical methods highlights the adverse effects of aging on the fundamental capacities to encode the timing of sound onsets in continuous streams and suppress cortical responses to predictable stimuli. This evidence motivates future research on the link between basic timing functions and general cognition, across the lifespan.

## Author contributions

S.A.K., M.S., conceptualized the study. S.A.K., M.S., collected the data.

A.C. designed and performed data analyses.

A.C., S.A.K., M.S., L.B. interpreted the results.

A.C. wrote the first draft of manuscript.

All authors contributed to, revised, and approved the final version of the manuscript.

## Acknowledgments

We thank Ina Koch at the Max Planck Institute for Human Cognitive and Brain Sciences, Leipzig Germany for her support in data collection, and Dr. Christian Obermeier for his input on the experimental design and support in data collection.

