## Supplementary Materials for "Aging impacts basic auditory and timing processes" (from another version of this preprint: version 1, 2024-03-27)

|  | **SumSq** | **F** | **p** |
| --- | --- | --- | --- |
| **Intercept** | 81,62 | 6,681 | 0,005 |
| **Group * Time** | 7,78 | 0,637 | 0,496 |
| **Error** | 488,70 | 1 | 0,5 |

Table 1 – Repeated-measures ANOVA on the N100 peak amplitude.
The Repeated-measures ANOVA on the N100 peak amplitude tested the Group * Time interaction effect. Columns provide, in order, the Sum of squares, F-value and the p-value.

|  |  |  | **p** | **Obs Diff** | **Eff Size** |
| --- | --- | --- | --- | --- | --- |
| **P50** | **Amplitude** | Mean | 0,002 | 0,974 | 0,466 |
|  |  | Peak | 0,086 | 0,536 | 0,255 |
|  |  | Peak Latency | 0,009 | 0,006 | 0,451 |
|  | **Variance** | Mean | 0,001 | -163,441 | -0,255 |
|  |  | Peak | 0,001 | -181,825 | -0,263 |
|  |  | Peak Latency | 0,001 | -181,825 | -0,263 |
| **N100** | **Amplitude** | Mean | 0,001 | 2,238 | 0,832 |
|  |  | Peak | 0,001 | 2,792 | 0,905 |
|  |  | Peak Latency | 0,017 | 0,006 | 0,334 |
|  | **Variance** | Mean | 0,001 | -176,364 | -0,257 |
|  |  | Peak | 0,001 | -180,803 | -0,262 |
|  |  | Peak Latency | 0,001 | -180,803 | -0,262 |
| **P200** | **Amplitude** | Mean | 0,031 | 0,811 | 0,253 |
|  |  | Peak | 0,214 | 0,433 | 0,151 |
|  |  | Peak Latency | 0,283 | -0,003 | -0,149 |
|  | **Variance** | Mean | 0,001 | -149,294 | -0,256 |
|  |  | Peak | 0,001 | -171,271 | -0,267 |
|  |  | Peak Latency | 0,001 | -171,271 | -0,267 |

Table 2 – Group statistics on the ERP metrics.
The table reports the group statistics for ERP metrics across three ERP components: P50, N100, P200 (left column). For each ERP component, we assessed amplitude and variance (2^nd^ left column) in the mean amplitude, peak amplitude and peak latency in pre-defined time-windows of interest. From right to left, we report the effect size, observed difference and p values for the group comparisons as assessed by 1000 permutation testing.

|  | **p** | | | **Obs Diff** | | **Eff Size** |
| --- | --- | --- | --- | --- | --- | --- |
| **Oscillatory component at Sf** | | 0,004 | -1,637 | | -0,817 | |
| **Fractal component** | | 0,001 | -0,268 | |  | |
| **ITPC at Sf** | | 0,007 | 0,151 | |  | |

Table 3 – Analyses on Spectral parametrization and ITPC
Group analyses on the spectral parametrization focused on individual peak amplitudes at the stimulation frequency (Sf; first row), on the amplitude of the fractal component across the frequency spectrum (second row), and the inter-trial phase coherence (ITPC) peak at the Sf. In order, the columns report the p-value, observed group difference and effect size.

|  | **P50** | | **N100** | | **P200** | |
| --- | --- | --- | --- | --- | --- | --- |
|  | Peak Amp Var | Peak Lat Var | Peak Amp Var | Peak Lat Var | Peak Amp Var | Peak Lat Var |
| **OSc** | 0,574 | 0,571 | 0,571 | 0,574 | 0,571 | 0,571 |

Table 4 – Correlation between Oscillatory component at Sf and ERP metrics.
The table reports rho values from the correlations between the Peak amplitude of the oscillatory component (OSc) at the stimulation frequency and ERP metrics of variability in Peak amplitude (Peak Amp Var) and Peak latency (Peak Lat Var) in the P50, N100 and P200 components of the ERPs.

|  | **SumSq** | **F** | **p** | **tStat** |
| --- | --- | --- | --- | --- |
| **P50 Peak Lat** | 0,123 | 7,716 | 0,009 | 2,778 |
| **P200 Peak Lat** | 0,018 | 1,112 | 0,300 | -1,055 |
| **N100 Amp Var** | 0,107 | 6,685 | 0,015 | -2,586 |
| **Error** | 0,494 | 1,000 | 0,500 |  |

Table 5 – Step-wise regression model between ITPC and ERP metrics.
The table reports the results from the step-wise regression model including the individual inter-trial phase coherence (ITPC) peak amplitudes at the stimulation frequency and ERP metrics of amplitude, latency and variability across the P50, N100 and P200 components of the ERPs. The resulting model featured as predictors the P50 peak latency, P200 peak latency and N100 peak amplitude variance (in order, from top to bottom on the left column). The table reports (column-wise) sum of squares, F-stats, p value and t-stats.
